# The Listening Effort Profile of Eye Movements: Easy, Difficult, and Impossible Speech Comprehension

**DOI:** 10.64898/2026.06.30.735702

**Authors:** Björn Herrmann, Lauren K. Fink, Priya R. Pandey, Ingrid S. Johnsrude, Jennifer D. Ryan

**Author notes:** Correspondence concerning this article should be addressed to Björn Herrmann, Rotman Research Institute, Baycrest Academy for Research and Education, 3560 Bathurst St, North York, ON, M6A 2E1, Canada.

## Abstract

Speech comprehension in noisy environments often requires cognitive effort, but listeners may disengage when comprehension becomes impossible. Eye movements have recently emerged as a promising new measure of listening effort, but it remains unclear whether eye movements are sensitive to the full effort profile across easy, difficult, and impossible speech comprehension. Across four experiments, participants listened to sentences at easy, difficult, and impossible levels of multi-talker background babble while pupil size and eye movements were recorded. Pupil size generally followed the expected inverted u-shaped effort profile: low for easy-to-understand speech, maximal for difficult but still intelligible speech and lower again for impossible-to-understand speech, although this pattern partly reflected sustained, condition-specific differences and not only sentence-evoked responses. Gaze dispersion – measuring the spread of eye movements – decreased specifically during the sentence-listening period for difficult relative to easy and impossible speech, indicating reduced eye movements during active, effortful listening. However, gaze dispersion was also lower, but less specific to the sentence-listening period, during impossible compared to easy listening, especially in non-baseline-corrected analyses, suggesting that reduced eye movements do not index listening effort uniquely. Instead, eye movements appear to reflect both attentional engagement during difficult listening and disengagement or inward attention when meaningful listening is no longer possible. These findings indicate that pupil size and eye movements provide complementary indices of listening-related cognition, and highlight the integration of listening, cognition, and motor systems.

## Introduction

Speech comprehension in everyday environments is often challenged by background sound, such as competing talkers in a restaurant. Under such conditions, listeners must rely more strongly on cognitive resources to extract meaning from the masked acoustic signal. This increased cognitive investment is commonly referred to as listening effort (Pichora-Fuller et al., 2016; Peelle, 2018; Herrmann and Johnsrude, 2020). Assessing listening effort is important because people may experience speech comprehension as effortful even when performance remains relatively high (Kadem et al., 2020; Winn and Teece, 2021), and such effort may provide an early indication of hearing-related difficulties before overt communication failures become apparent (Pichora-Fuller et al., 2016; Helfer and Jesse, 2021).

One of the most widely used objective measures of listening effort is pupillometry (Winn et al., 2018; Zekveld et al., 2018). Pupil size typically increases when speech comprehension becomes more demanding, for example when speech is masked by background noise or degraded acoustically (Zekveld et al., 2010; Zekveld and Kramer, 2014; Ohlenforst et al., 2017a; Ohlenforst et al., 2018; Wendt et al., 2018; Kadem et al., 2020; Neagu et al., 2023; Baldock et al., 2024). Pupillometry has therefore provided an important physiological window into the cognitive demands of listening. However, pupil size is also sensitive to factors unrelated to listening effort, including luminance, gaze position relative to the eye-tracking camera, and arousal-related fluctuations (Brisson et al., 2013; Knapen et al., 2016; Mathôt, 2018; Fink et al., 2024). Moreover, although effects of SNR on pupil size have been shown for continuous speech of about 30 s (Seifi Ala et al., 2020; Fiedler et al., 2021) and for 60-s audiobook passages (Johns et al., 2024; with effects depending on baseline pupil size), other work highlights that pupillometric effects can be less reliable in more naturalistic, continuous speech-listening paradigms than in trial-based sentence-listening tasks (Cui and Herrmann, 2023; Herrmann et al., 2025; see also Widmann et al., 2025).

A growing body of work suggests that eye movements may provide an alternative or complementary marker of listening effort. When speech listening becomes more difficult, listeners reduce their eye movements, as reflected in fewer micro-saccades and reduced spatial gaze dispersion (Contadini-Wright et al., 2023; Cui and Herrmann, 2023; Herrmann and Ryan, 2024; Cui et al., 2025; He et al., 2025; Herrmann et al., 2025; Mariotti et al., 2026). These reductions have been observed across different speech materials, including isolated sentences and continuous stories, across different visual contexts, including blank screens, fixation, and moving-dot displays, and across stationary and mobile devices (Cui and Herrmann, 2023; Herrmann and Ryan, 2024; Cui et al., 2025; Herrmann et al., 2025). Mechanistically, reduced eye movements during effortful listening may help limit distracting visual input or may reflect broader interactions between oculomotor, attentional, and auditory systems. For example, periods with reduced eye movements have been associated with changes in auditory cortical excitability that could support sound processing (Schneider and Mooney, 2015, 2018; O’Connell et al., 2020).

A useful physiological measure of listening effort should be sensitive not only to increases in task demand, but also to disengagement. Listeners do not always continue investing effort as listening becomes more difficult. When the demands of a listening situation exceed the listener’s available resources, or when the expected reward no longer justifies the required cognitive investment, listeners may disengage or “give up” (Yerkes and Dodson, 1908; Brehm and Self, 1989; Richter, 2013; Westbrook and Braver, 2015; Heffernan et al., 2016; Pichora-Fuller et al., 2016; Herrmann and Johnsrude, 2020). Thus, effort measures should capture the full effort profile: low effort when speech is easy, increased effort when speech is difficult but still potentially understandable, and reduced effort again when comprehension becomes impossible.

Pupillometry appears to follow this expected inverted u-shape effort profile in many listening studies. Pupil size is often largest when speech is difficult but still intelligible, and smaller when speech is either easy to understand or impossible to understand (Zekveld and Kramer, 2014; Ohlenforst et al., 2017b; Ohlenforst et al., 2018; Wendt et al., 2018; Herrmann and Ryan, 2024). Eye movements, however, may follow a different pattern. In previous work, gaze dispersion – a measure of the spread of a person’s eye movements over a period of time – decreased when speech comprehension was difficult compared to easy, but also remained reduced when speech comprehension was impossible (Herrmann and Ryan, 2024). This sustained reduction was observed even after sentence offset, when acoustic stimulation no longer differed between easy, difficult, and impossible conditions, and even when participants could anticipate that an upcoming trial would be impossible. These findings suggest that pupil size and eye movements may index different aspects of listening-related cognition (Herrmann and Ryan, 2024; Liu and Chait, 2026).

Why eye movements remain low during impossible listening is unresolved. One possibility is that visual stimulation in previous work – a moving-dot display – encouraged listeners to suppress visual exploration, thereby producing reduced eye movements that were not specific to listening (Herrmann and Ryan, 2024). Another possibility is that listeners continued searching for a sentence in the background noise, even when comprehension was impossible. Because the sentence was inaudible, listeners may not have known when it ended and may have continued orienting attention to the auditory stream after the actual sentence offset. According to this possibility, eye movements should eventually increase if the post-sentence interval is long enough for listeners to infer that no more speech will occur. A third possibility is that reduced eye movements during impossible listening reflect a different state altogether, such as disengagement or mind wandering, since inward attention can reduce eye movements (Reichle et al., 2010; Benedek et al., 2017; Walcher et al., 2017; Mézière et al., 2026). If so, reduced eye movements might persist throughout a listening trial, even before or after the time window in which speech could be processed, especially when the visual scene provides little reason to explore.

The current study investigated these possibilities across four experiments by examining pupil size and gaze dispersion during easy, difficult, and impossible speech comprehension. In Experiment 1, we tested whether reduced gaze dispersion during impossible listening occurs in the absence of any visual stimulation, using a blank screen rather than a moving-dot display. In Experiment 2, we extended the post-sentence interval to determine whether reduced eye movements in the impossible condition would recover once listeners had more time to realize that the sentence had ended. In Experiment 3, we removed the sentence entirely from the impossible condition and informed participants that no sentence would be present, thereby further discouraging continued attempts to extract speech from the noise. In Experiment 4, we tested whether visually engaging stimulation would increase eye movements during periods of disengagement and reveal a more selective reduction during difficult speech processing. Together, these experiments provide a comprehensive assessment of how pupil size and eye movements track the transitions from easy listening, to effortful listening, to impossible or disengaged listening.

## General methods

### Participants

Participants were native English speakers or individuals who learned English before the age of 5 years. They reported having normal hearing and neither had a prescription for hearing aids or wore them. Participants gave written informed consent prior to the experiment and were paid $10 CAD per half-hour for their participation. The study was conducted in accordance with the Declaration of Helsinki, the Canadian Tri-Council Policy Statement on Ethical Conduct for Research Involving Humans (TCPS2-2014), and was approved by the Research Ethics Board of the Rotman Research Institute at Baycrest. Detailed demographic information about participants is provided in the individual experiment sections below.

### Experimental setup

Participants sat in a single-wall sound booth (Eckel). Experimental procedures were run using Psychtoolbox (v3.0.14) in MATLAB (MathWorks Inc.) on a Lenovo T450s laptop with Microsoft Windows 7 (Experiments 1-3) or a desktop computer with Windows 10 (Experiment 4), all in the same space. The laptop/computer screen was mirrored to a ViewSonic monitor in the sound booth. Participants sat in front of the ViewSonic (60 Hz refresh rate) monitor at a distance of about 70 cm with their head stabilized through a chin and forehead rest. Sounds were presented using a RME Fireface UCX external sound card and Sony Dynamic Stereo MDR-7506 headphones at a comfortable listening level of ∼70 dB SPL.

### Stimulus materials and procedure

Stimuli comprised auditory sentences from the Harvard sentence lists (male native English speaker; average duration: ∼2.5 seconds) (IEEE, 1969) obtained from a publicly accessible resource (speaker ID: ‘PNM055’; https://depts.washington.edu/phonlab/projects/uwnu.php; Panfili et al., 2017; McCloy et al., 2018). A new set of sentences was selected for each experiment. Each sentence was embedded within a 12-talker babble background noise (Bilger, 1984), with the sentence beginning 2.8 s after the babble onset (Experiment 1; similar to Herrmann and Ryan, 2024) or 1.5 s after babble onset (Experiments 2-4). The duration of the babble masker was 6.3 s in Experiment 1 and 8.5 s in Experiments 2-4. All sentences terminated before the babble ended. Sentences were presented at three signal-to-noise ratios (SNRs): +11 dB (easy), -4 dB (difficult), and -19 dB (impossible). In Experiments 3 and 4, the impossible condition did not contain any sentence (see below). Listening effort was expected to be minimal at the extreme SNRs (easy and impossible), as participants either required little cognitive effort or were unable to understand the sentence and presumably would stop trying. In contrast, the highest effort was anticipated in the difficult, but still intelligible, SNR condition. Different SNRs were created by modifying the level of sentences while keeping the level of the babble noise constant across trials (Ohlenforst et al., 2017b; Kadem et al., 2020; Cui and Herrmann, 2023; Herrmann and Ryan, 2024). This ensured that stimuli with different SNRs did not differ prior to sentence onset and after sentence offset (Ohlenforst et al., 2017b; Kadem et al., 2020; Cui and Herrmann, 2023).

The computer screen was blank and mid-gray before and during sound stimulation (except in Experiment 4). Participants were free to look around the screen however they wished (but keep their gaze on the screen), while reducing blinks (Johansson et al., 2006; Johansson et al., 2011; Johansson et al., 2012; Cui and Herrmann, 2023; Herrmann and Ryan, 2024). A probe word appeared on the screen after the sound on each trial, and participants had to decide whether it was semantically related or unrelated to the preceding sentence with a keypress (Rodd et al., 2010a; Rodd et al., 2010b; Kadem et al., 2020; Cui and Herrmann, 2023). Participants were asked to guess if they did not know.

Each SNR condition included 60 trials, distributed across five blocks. Within each block, participants listened to 12 sentences at each SNR. Trials were grouped by SNR, meaning 12 sentences with the same SNR were presented consecutively. Participants were informed in advance of the upcoming listening difficulty (easy, difficult, or impossible) to enable them to prepare for the upcoming trials (Herrmann and Ryan, 2024). Had the SNR conditions been randomized on a trial-by-trial basis, participants would not have known when a sentence would be unintelligible, and thus might have attempted to exert effort even in the impossible condition – an outcome that would have confounded our investigation.

To prevent potential confounding of specific sentences with SNR conditions, the sentence-to-SNR mapping was randomized across participants. Prior to the main experiment, a brief training block of 12 trials was used to familiarize participants with the semantic-relatedness task and the eye-tracking calibration process.

### Behavioral data analysis

For each SNR condition, the proportion of correct responses in the semantic-relatedness task was calculated for display. Statistical analyses were carried out using single-trial, binary responses within a generalized linear mixed-effects model with a binomial distribution and logit link function (MATLAB; fitglme, Statistics and Machine Learning Toolbox), fit by Laplace approximation. The model included the fixed effect of SNR (+11 [easy], -4 [difficult], -19 dB or no sentence [impossible]) and random intercepts and random SNR slopes for participants and for sentences: data ∼ 1 + SNR + (1 + SNR | participant) + (1 + SNR | sentence). Effects coding was used, such that the omnibus effect of SNR was assessed with an F test. Differences between SNR conditions were examined using pairwise contrasts of the fixed effects, for which we report the coefficient (b; in log odds) with its 95% confidence interval, the corresponding odds ratio, the t value with residual degrees of freedom, and p values corrected across contrasts using Holm’s method (Holm, 1979). Performance in the impossible condition was compared against chance level (50%) by testing whether the model-estimated log odds of a correct response in that condition differed from zero.

### Pupillometry and eye-movement recordings

The pupil size and eye movements were concurrently recorded from the left and right eye using an EyeLink Duo eye-tracker (SR Research Ltd.) at a sampling rate of 500 Hz. In the beginning of each experimental block, a nine-point fixation was used to calibrate the eye-tracker (McIntire et al., 2014).

### Processing of pupil-size and eye-movement data

Data analysis was conducted in MATLAB software (MathWorks Inc.). The main metrics were pupil size and gaze dispersion. Gaze dispersion is a measure of spread or variability of eye movements over time that non-specifically captures any eye movement changes associated with difficult listening (Cui and Herrmann, 2023; Herrmann and Ryan, 2024; Cui et al., 2025; Herrmann et al., 2025). Gaze dispersion behavior in response to listening challenge is similar to that of (micro-)saccade rate (Contadini-Wright et al., 2023; Herrmann et al., 2025). However, gaze dispersion is preferred here because it appears to be more sensitive to listening challenges than (micro-)saccade rate (Herrmann et al., 2025), likely in part because (micro-)saccade data are sparse (occurring ∼1-2 per second), whereas gaze dispersion is based on the continuous time series data.

Preprocessing was calculated separately for each block and separately for each eye. Missing data points in the pupil and x–y eye-coordinate time courses resulting from blinks or other artifacts were set to NaN (‘not a number’ in MATLAB), as were data points from 0.1 s before to 0.2 s after the artifact. Pupil-size values that differed from the mean pupil size by more than 3 times the standard deviation in a block were classified as outliers. The corresponding pupil-size, x, and y data points were set to NaN.

For the pupil size, NaN-coded data points were interpolated using MATLAB’s ‘pchip’ method. Pupil-size time courses were filtered with a 5-Hz low-pass filter (51 points, Kaiser window; β = 4). Pupil-size time courses for the left and right eye were averaged.

For gaze dispersion, NaN-coded points were ignored. Gaze dispersion was calculated as the standard deviation in gaze across time points, averaged across x- and y-coordinates, and transformed to logarithmic values to make the metric’s distributional properties more Gaussian. Smaller values indicate less gaze dispersion (i.e., fewer eye movements). To obtain time courses, gaze dispersion was calculated for 1-s (500-sample) sliding time windows centered sequentially on each time point. If more than 90% of data were unavailable within a 1-s time window (that is, 450 or more samples were NaN-coded), gaze dispersion for the corresponding time point was set to NaN and ignored during averaging (Cui and Herrmann, 2023; Herrmann and Ryan, 2024; Cui et al., 2025; Herrmann et al., 2025). Gaze-dispersion time courses for the left and right eye were averaged.

Continuous pupil-size and gaze-dispersion data were divided into single-trial time courses ranging from -1 to 6 s for Experiment 1 and -1 to 8.2 s for Experiments 2-4 time-locked to babble onset. Data for an entire trial were excluded from analysis if the percentage of NaN data entries made up more than 50% of the trial. For visualization, pupil size and gaze dispersion were averaged across trials, separately for each SNR condition.

Pupil size and gaze dispersion were analyzed with and without baseline correction. Non-baseline-corrected data enable examination of preparatory, sustained effects, whereas baseline-corrected data enable examination of trial-evoked responses. For the baseline-correction analysis, data were normalized relative to the -1 to 0 s prior to the onset of the babble noise by subtracting the mean baseline data from the data at each time point.

For statistical analyses, gaze dispersion was averaged across the 2.8–5.3-s (Experiment 1) or 1.5–4-s (Experiments 2-4) time window during which sentences were presented. Mean pupil size was calculated for a time window that was delayed by 0.5 s to 3.3–5.8-s for Experiment 1 and 2–4.5-s for Experiments 2-4, because the pupil size is known to change relatively slowly and peak late during sentence listening (Knapen et al., 2016; Winn and Moore, 2018; Winn et al., 2018; Kadem et al., 2020; Zhang et al., 2022). Data were also analyzed for two other time windows in order to investigate differences in eye metrics during periods when no active listening engagement was required to solve the main speech-comprehension task. These included the –1 to 0 s baseline time window (all experiments; non-baseline-corrected data) and a post-sentence time window (6–8 s) for Experiments 2-4.

Statistical analyses of pupil size and gaze dispersion were carried out using linear mixed-effects models (LMEs) in MATLAB (MathWorks Inc.; fitlme, Statistics and Machine Learning Toolbox), calculated on single-trial data. Separate models were calculated for each experiment, each measure (pupil size, gaze dispersion), and for baseline-corrected and non-baseline-corrected data. Each model included the fixed effect of SNR (easy, difficult, impossible) and, when more than one time window was analyzed, the fixed effect of time window and the SNR × time window interaction, together with random intercepts and random SNR slopes for participants and for sentences: data ∼ 1 + SNR × time window + (1 + SNR | participant) + (1 + SNR | sentence). Models with the full SNR × time window random-effects structure did not converge and were therefore reduced to the structure above. Omnibus effects were assessed with F tests using effects coding, with partial eta squared calculated as η_p_² = (F · df₁) / (F · df₁ + df₂). Differences between SNR conditions were examined using pairwise contrasts, separately for each time window, by re-parameterizing the model so that the respective SNR level served as the reference level. The corresponding fixed-effect coefficient (b) reflects the difference between the two conditions in the units of the dependent measure. We report b with its (uncorrected) 95% confidence interval, the t value with residual degrees of freedom, and p values corrected across all contrasts using Holm’s method (Holm, 1979).

## Experiment 1

Experiment 1 aimed to replicate the finding that listeners show reduced eye movements during impossible-to-understand speech compared to easy-to-understand speech (Herrmann and Ryan, 2024), but in the absence of any visual stimulation. This experiment aimed to establish whether the previously observed reduction in eye movements was influenced by the concurrent visual input used in our earlier work (Herrmann and Ryan, 2024), or whether the effect also occurs when no visual stimuli are present.

### Methods and materials

#### Participants

Sample sizes were based on our previous work (Cui and Herrmann, 2023; Herrmann and Ryan, 2024). Twenty-four adults participated in Experiment 1 (median age: 25 years, age range: 18–35 years; 11 male, 12 female, 1 transgender male). Data from one additional participant were recorded but not analyzed because their proportion of correct responses for the easiest condition in the behavioral task was equal to or lower than 0.6, indicating low task compliance. Eight individuals also participated in one other experiment, one in two other experiments, and another person in the three other experiments (sessions were several weeks apart).

#### Stimulation and procedures

Stimulation and procedures were as described above. Participants listened to speech in babble at easy, difficult, and impossible SNRs while the computer screen was blank. Information provided to participants was “Comprehension of the next 12 sentences will likely be: “ followed by “easy”, “difficult”, or “impossible”. They performed a semantic-relatedness task to test for sentence comprehension. Pupil size and gaze dispersion were calculated for epochs ranging from -1 s to 6 s, time-locked to noise onset.

### Results

#### Behavior

Performance in the semantic relatedness task was higher for: 1) easy than difficult, 2) easy than impossible, and 3) difficult than impossible SNRs (mean accuracy: 0.921, 0.763, and 0.499 for easy, difficult, and impossible, respectively; for all b ≥ 1.32, 95% CI [≥1.05, ≥1.59], OR ≥ 3.76, t_4317_ ≥ 9.60, p_Holm_ < 0.001; effect of SNR: F_2,4317_ = 109.01, p = 6.5 · 10^−47^). Performance in the impossible condition did not differ from chance level (b = -0.007, 95% CI [-0.121, 0.107], t_4317_ = - 0.12, p = 0.902).

#### Pupil size

For non-baseline-corrected data, SNR effects differed between different time windows (SNR × time window interaction: F_2,7638_ = 8.022, p = 3.3 · 10^−4^, η_p_^2^ = 0.002). For the sentence-response time window (3.3 to 5.8 s), the pupil size was larger for the difficult than for the impossible condition (b = 40.44, 95% CI [21.52, 59.36], t_7638_ = 4.19, p_Holm_ = 2.8 · 10^−5^) and the easy condition (b = 50.226, 95% CI [29.93, 70.52], t_7638_ = 4.852, p_Holm_ = 1.2 · 10^−6^; Figure 1A). No significant differences after multiple-comparison correction were found for the pre-noise time window (-1 to 0 s; for all p_Holm_ > 0.1).

**Figure 1:**
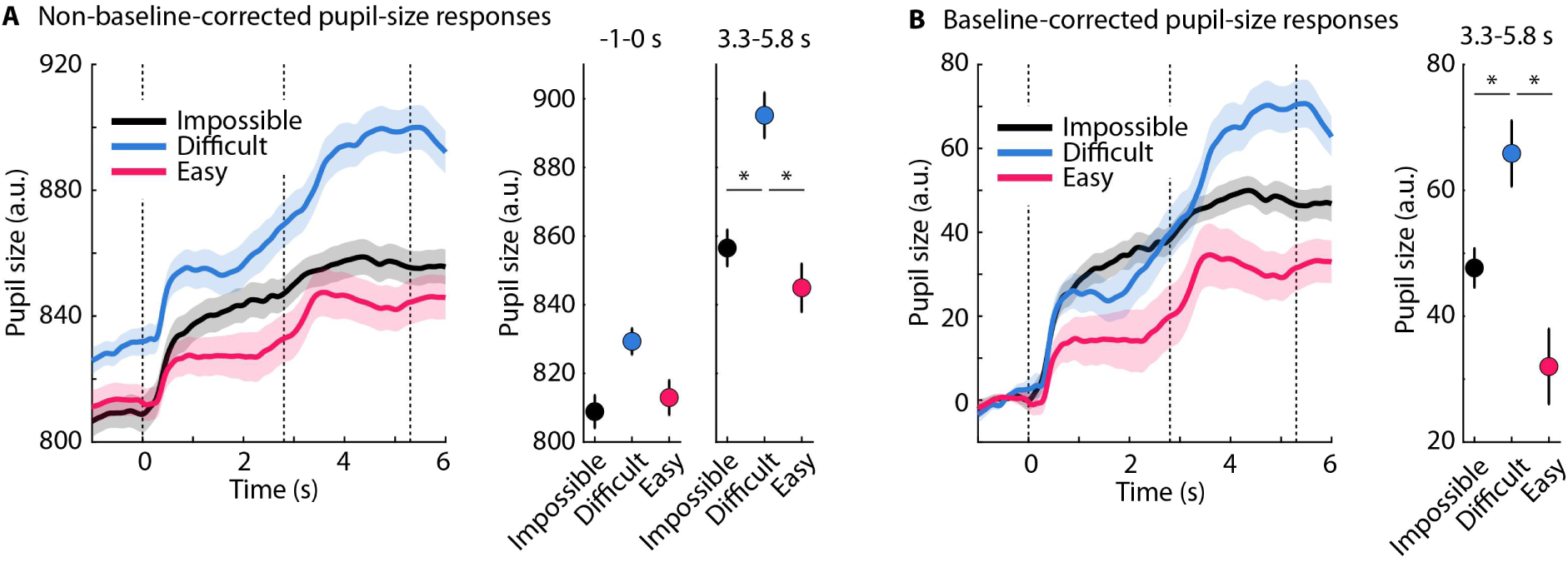
Pupil size results for Experiment 1. **A:** Non-baseline-corrected data, with time-courses shown on the left and mean pupil size for two time windows shown on the right. The dashed, vertical lines in the time-course plot mark the babble onset, sentence onset, and average sentence offset, from left to right respectively. Shaded areas around the time-courses and error bars reflect the standard error of the mean after removal of between-participant variance. *p_Holm_ ≤ 0.05; only significant effects are indicated. **B:** Same as for Panel A, for baseline-corrected pupil size.

For the baseline-corrected data in the 3.3 to 5.8 s time window, the LME model showed a larger pupil size for the difficult compared to the impossible condition (b = 19.06, 95% CI [5.27, 32.84], t_3819_ = 2.71, p_Holm_ = 0.014) and the easy condition (b = 34.54, 95% CI [13.63, 55.44], t_3819_ = 3.24, p_Holm_ = 0.004; effect of SNR: F_2,7638_ = 5.702, p = 0.003, η_p_^2^ = 0.003; Figure 1B).

#### Gaze dispersion

SNR effects differed between different time windows (SNR × time window interaction for non-baseline-corrected and baseline-corrected data: F > 11, p = 1 · 10^−5^, η_p_^2^ > 0.006). There were no differences between SNR conditions in the pre-noise time window (-1 to 0 s; p_Holm_ = 1). However, for both the non-baseline-corrected data and the baseline-corrected data, gaze dispersion was lower for the difficult condition compared to both the impossible condition (b ≥ 0.06, 95% CI [≥0.02, ≥0.10], t > 2.5, p_Holm_ < 0.04) and the easy condition (b ≥ 0.13, 95% CI [≥0.08, ≥0.17], t > 4.5, p_Holm_ < 0.001), and lower for the impossible relative to the easy condition (b ≥ 0.07, 95% CI [≥0.03, ≥0.11], t > 3.1, p_Holm_ < 0.005; Figure 2).

**Figure 2:**
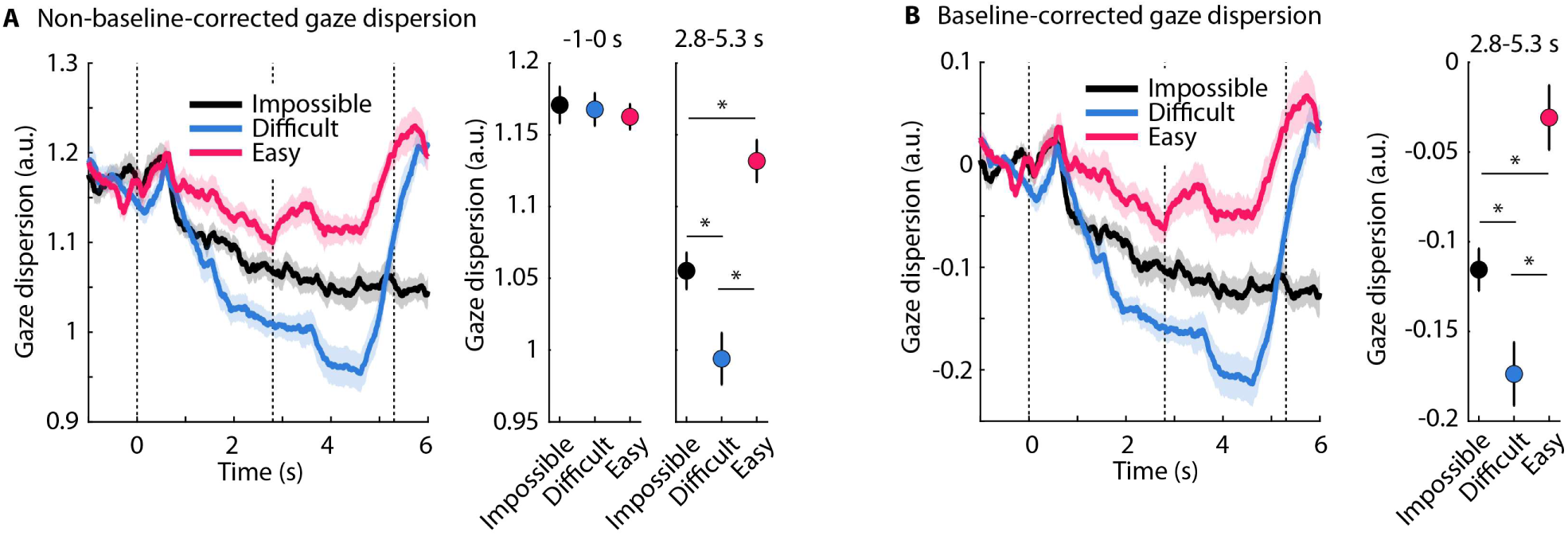
Gaze dispersion results for Experiment 1. **A:** Non-baseline-corrected data, with time-courses shown on the left and mean gaze dispersion for two time windows shown on the right. The dashed, vertical lines in the time-course plot mark the babble onset, sentence onset, and average sentence offset, from left to right respectively. Shaded areas around the time-courses and error bars reflect the standard error of the mean after removal of between-participant variance. *p_Holm_ ≤ 0.05; only significant effects are indicated. **B:** Same as for Panel A, for baseline-corrected gaze dispersion.

#### Summary

Experiment 1 showed the expected listening-effort profile for pupil size, with larger pupil responses for difficult speech than for easy and impossible speech (Zekveld and Kramer, 2014; Ohlenforst et al., 2018; Herrmann and Ryan, 2024). Gaze dispersion was lower for difficult compared with easy speech, consistent with reduced eye movements during effortful listening (Contadini-Wright et al., 2023; Cui et al., 2025). Critically, gaze dispersion was also reduced for impossible compared with easy speech even at the very end of a trial, replicating and extending prior work in which participants viewed a moving-dot display during listening (Herrmann and Ryan, 2024). The present experiment shows that this reduction occurs even when the screen is blank. Thus, reduced eye movements during listening to speech that is impossible to understand cannot be attributed to suppression of visual exploration caused by concurrent visual stimulation. Instead, the effect appears to arise independently of visual input, suggesting that reduced gaze dispersion in the impossible condition may reflect some cognitive process rather than a response to the visual environment.

## Experiment 2

In Experiment 2, we tested whether reduced eye movements in the impossible condition reflect continued attempts to identify a sentence in noise. Even when speech comprehension is impossible, participants may still listen for a sentence and attempt to extract it from the babble, and this continued search could reduce eye movements. Because the sentence is not perceptible in the impossible condition, participants also cannot determine when it ends. As a result, they may continue searching for the sentence even after its actual offset, which could explain why gaze dispersion remains reduced after sentence presentation has ended. To test this possibility, Experiment 2 extended the duration of the babble noise after sentence offset. If listeners were continuing to search for the sentence, then the longer post-sentence interval should allow them to realize, based on the timing of sentences in the easy and difficult conditions, that no further speech is likely to occur. Listeners should eventually give up searching, and eye movements should increase during the extended post-sentence period.

### Methods and materials

#### Participants

Twenty-five adults participated in Experiment 2 (median age: 24 years, age range: 19–35 years; 12 male, 11 female, 1 non-binary, 1 transgender male). Data from three additional participants were recorded but not analyzed because their proportion of correct responses for the easiest condition in the behavioral task was lower than 0.6 (n=1) or their physiological recordings contained more than 50% of missing data on more than 50% of trials (n=2). Ten individuals also participated in one other experiment, one individual in two other experiments, and another person in the three other experiments (sessions were several weeks apart).

#### Stimulation and procedures

Stimulation and procedures were as described above. Participants listened to a new set of sentences in babble at easy (+11 dB), difficult (-4 dB), and impossible (-19 dB) SNRs while the computer screen was blank. The babble played for 8.5 s, with the sentence starting 1.5 s after babble onset and the average sentence offset at around 4 s. Experiment 2 therefore differed from Experiment 1 in two respects: the babble continued for longer after sentence offset (4.5 s vs 1 s on average), and the sentence began earlier relative to babble onset (1.5 s vs 2.8 s). The earlier onset compensated for the longer babble, keeping overall trial duration short enough to preserve the same number of trials. Participants performed the semantic-relatedness task to test for sentence comprehension. Pupil size and gaze dispersion were calculated for epochs ranging from -1 s to 8.2 s, time-locked to noise onset (note that the final 0.3 s were excluded, to avoid capturing smear-back effects from the response screen following babble offset due to the 1-s sliding window used for the gaze dispersion analysis).

### Results

#### Behavior

Performance in the semantic relatedness task was higher for the easy condition than for both the difficult and impossible conditions, and higher for the difficult than for the impossible condition (mean accuracy: 0.897, 0.736, and 0.519 for easy, difficult, and impossible, respectively; for all b ≥ 1.03, 95% CI [≥0.82, ≥1.24], OR ≥ 2.81, t_4461_ ≥ 9.57, p_Holm_ < 0.001; effect of SNR: F_2,4461_ = 131.52, p =3.2 · 10^−56^). Performance in the impossible condition did not differ from chance level (b = 0.077, 95% CI [-0.036, 0.190], t_4461_ = 1.33, p = 0.183).

#### Pupil size

For non-baseline-corrected data, SNR effects differed between time windows (SNR × time window interaction: F_4,11709_ = 8.917, p = 3.5 · 10^−7^, η_p_^2^ = 0.003). In the sentence-response time window (2 to 4.5 s), pupil size was greater in the difficult than in the impossible condition (b = 33.70, 95% CI [13.06, 54.35], t_11709_ = 3.20, p_Holm_ = 0.012) and the easy condition (b = 31.11, 95% CI [9.51, 52.72], t_11709_ = 2.82, p_Holm_ = 0.038; Figure 3A). Other contrasts did not significantly differ after multiple-comparison correction (all p_Holm_ > 0.05).

**Figure 3:**
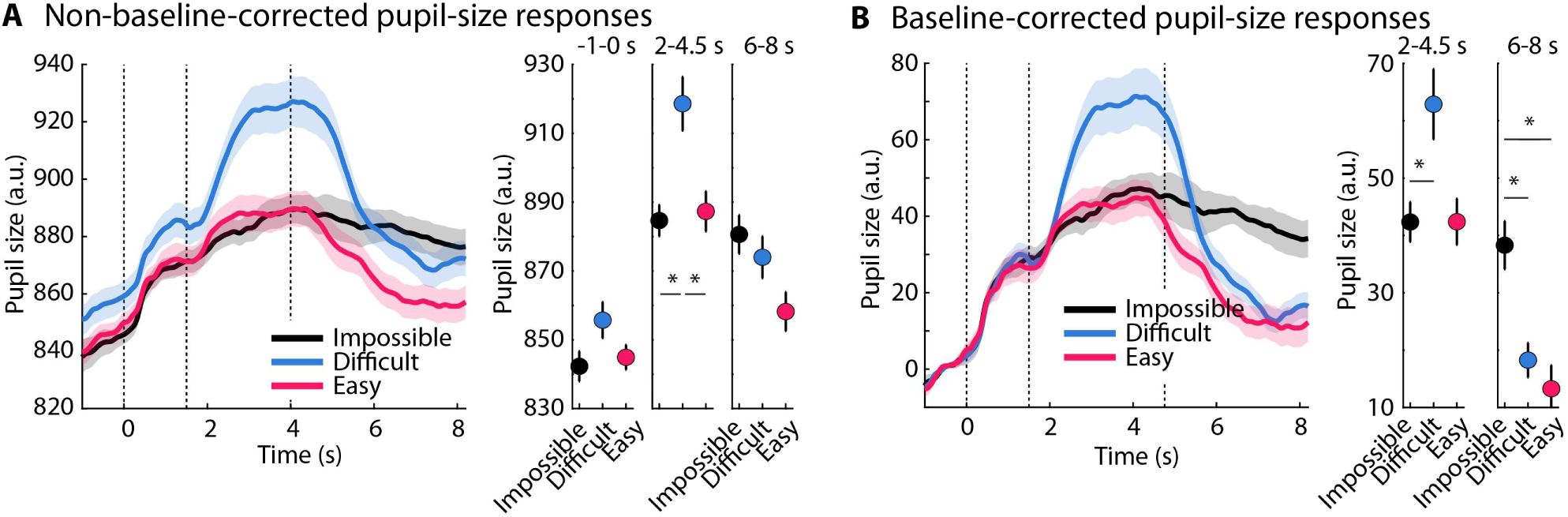
Pupil size results for Experiment 2. **A:** Non-baseline-corrected data, with time-courses shown on the left and mean pupil size for three time windows shown on the right. The dashed, vertical lines in the time-course plot mark the babble onset, sentence onset, and average sentence offset, from left to right respectively. Shaded areas around the time-courses and error bars reflect the standard error of the mean after removal of between-participant variance. *p_Holm_ ≤ 0.05; only significant effects are indicated. **B:** Same as for Panel A, for baseline-corrected pupil size.

For baseline-corrected data, the SNR × time window interaction was significant (F_2,7806_ = 14.519, p = 5.1 · 10^−7^, η_p_^2^ = 0.004). In the sentence-response time window (2 to 4.5 s), the pupil size was greater in the difficult condition than in the impossible (b = 20.26, 95% CI [4.75, 35.77], t_7806_ = 2.56, p_Holm_ = 0.042), whereas the difficult vs easy contrast was not significant after multiple-comparison correction (b = 19.07, 95% CI [2.86, 35.28], t_7806_ = 2.31, p_Holm_ = 0.063). In the post-sentence time window (6 to 8 s), pupil size was greater in the impossible than in both the difficult condition (b = 20.86, 95% CI [5.35, 36.37], t_7806_ = 2.64, p_Holm_ = 0.042) and the easy condition (b = 24.33, 95% CI [10.36, 38.30], t_7806_ = 3.41, p_Holm_ = 0.004; Figure 3B).

#### Gaze dispersion

For non-baseline-corrected gaze dispersion, the SNR × time window interaction was significant (F_4,11596_ = 40.703, p = 6.3 · 10^−34^, η_p_^2^ = 0.014). In the pre-noise time window (-1 to 0 s), there was less gaze dispersion for the impossible condition than the difficult condition (b = 0.07, 95% CI [0.03, 0.12], t_11596_ = 3.39, p_Holm_ = 0.003). In the sentence-response time window (1.5 to 4 s), gaze dispersion was smaller in both the impossible condition (b = 0.09, 95% CI [0.04, 0.13], t_11596_ = 3.77, p_Holm_ = 0.001) and the difficult condition (b = 0.13, 95% CI [0.09, 0.17], t_11596_ = 5.94, p_Holm_ = 2.1 · 10^−8^) relative to the easy condition. In the post-sentence time window (6 to 8 s), there was less gaze dispersion in the impossible relative to the difficult (b = 0.17, 95% CI [0.13, 0.21], t_11596_ = 7.97, p_Holm_ = 1.6 · 10^−14^) and the easy condition (b = 0.15, 95% CI [0.10, 0.19], t_11596_ = 6.41, p_Holm_ = 1.2 · 10^−9^; Figure 4A).

**Figure 4:**
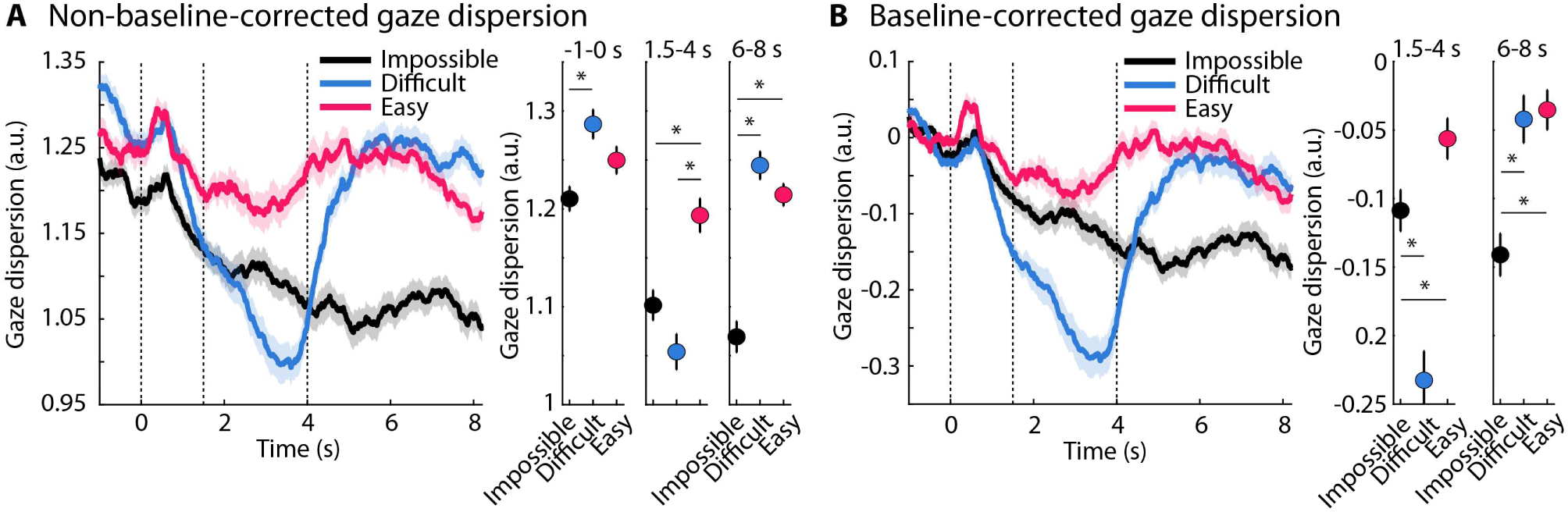
Gaze dispersion results for Experiment 2. **A:** Non-baseline-corrected data, with time-courses shown on the left and mean gaze dispersion for three time windows shown on the right. The dashed, vertical lines in the time-course plot mark the babble onset, sentence onset, and average sentence offset, from left to right respectively. Shaded areas around the time-courses and error bars reflect the standard error of the mean after removal of between-participant variance. *p_Holm_ ≤ 0.05; only significant effects are indicated. **B:** Same as for Panel A, for baseline-corrected gaze dispersion.

The results for baseline-corrected data are mostly similar to those for non-baseline corrected data (SNR × time window interaction: F_2,7592_ = 37.759, p = 4.8 · 10^−17^, η_p_^2^ = 0.010). In the sentence-response time window, gaze dispersion was lower for the difficult condition compared to both the easy condition (b = 0.16, 95% CI [0.10, 0.22], t_7592_ = 5.36, p_Holm_ = 5.3 · 10^−7^) and the impossible condition (b = 0.12, 95% CI [0.06, 0.18], t_7592_ = 3.86, p_Holm_ = 4.6 · 10^−4^). In the post-sentence time window, gaze dispersion was lower in the impossible than in the other two conditions (difficult: b = 0.10, 95% CI [0.04, 0.16], t_7592_ = 3.30, p_Holm_ = 0.003; easy: b = 0.11, 95% CI [0.06, 0.16], t_7592_ = 4.17, p_Holm_ = 1.5 · 10^−4^; Figure 4B).

#### Summary

As in Experiment 1, pupil size showed the expected effort-related pattern during sentence processing, with larger pupil responses for difficult than easy or impossible speech. Interestingly, pupil size also remained elevated for the impossible condition after sentence offset, especially in the baseline-corrected data, suggesting a trial-evoked response that persisted into the extended post-sentence interval. Gaze dispersion was reduced for difficult compared with easy speech, again indicating reduced eye movements during effortful listening. Critically, however, extending the post-sentence interval did not lead to a recovery of eye movements in the impossible condition. Instead, gaze dispersion remained lower for impossible speech than for easy speech across time windows. Thus, reduced eye movements in the impossible condition do not appear to reflect a temporary search for a sentence that resolves once listeners have enough time to realize that the sentence has ended. Rather, the sustained reduction suggests that impossible listening may involve a more persistent cognitive state beyond the period of sentence presentation, potentially reflecting continued auditory orienting or a shift toward disengagement-related processes that Experiment 3 was designed to examine more directly.

## Experiment 3

In Experiments 1 and 2, participants knew that they were unlikely to understand speech in the impossible condition. Nevertheless, because a sentence was still present, they may have attempted to understand it anyway. Simply knowing that a sentence existed may have prompted participants to listen attentively, which could have contributed to the reduced eye movements observed in the impossible condition. To further encourage giving up attentive listening, Experiment 3 followed the same procedures as Experiment 2 but did not present a sentence in the babble during the impossible condition and explicitly informed participants that no sentence was present.

### Methods and materials

#### Participants

Twenty-six adults participated in Experiment 3 (median age: 22 years, age range: 19–33 years; 4 male, 21 female, 1 non-binary). Data from four additional participants were recorded but not analyzed because more than 50% of data were missing from physiological recordings on more than 50% of trials. Ten individuals also participated in one other experiment, three people in two other experiments, and another person in the three other experiments (sessions were several weeks apart).

#### Stimulation and procedures

Experimental procedures were identical to those in Experiment 2, with the exception that no sentence was presented in the ‘impossible’ condition. Participants were informed that no sentence was present during the impossible condition, using the following wording: “No sentence, only noise is presented for the next 12 trials. Just randomly select ‘related’ or ‘unrelated’ when asked.”.

### Results

#### Behavior

Performance in the semantic-relatedness task was higher in the easy condition than in either the difficult or impossible conditions, and higher in the difficult condition compared to the impossible condition (mean performance: 0.912, 0.762, and 0.496 for easy, difficult, and impossible, respectively; for all b ≥ 1.33, 95% CI [≥1.08, ≥1.58], OR ≥ 3.78, t_4677_ ≥ 10.55, p_Holm_ < 0.001; effect of SNR: F_2,4677_ = 129.99, p = 1.2 · 10^−55^). Performance in the impossible condition did not differ from chance level (b = -0.018, 95% CI [-0.131, 0.095], t_4677_ = -0.31, p = 0.755).

#### Pupil size

Non-baseline corrected data showed different SNR effects across time windows (interaction: F_4,12366_ = 8.485, p = 7.8 · 10^−7^, η_p_^2^ = 0.002). In the sentence-response time window, pupil size was larger in the difficult condition than in either the impossible (b = 33.89, 95% CI [12.63, 55.15], t_12366_ = 3.12, p_Holm_ = 0.014) or the easy condition (b = 31.29, 95% CI [14.23, 48.35], t_12366_ = 3.59, p_Holm_ = 0.003; Figure 5A). No other differences were found.

**Figure 5:**
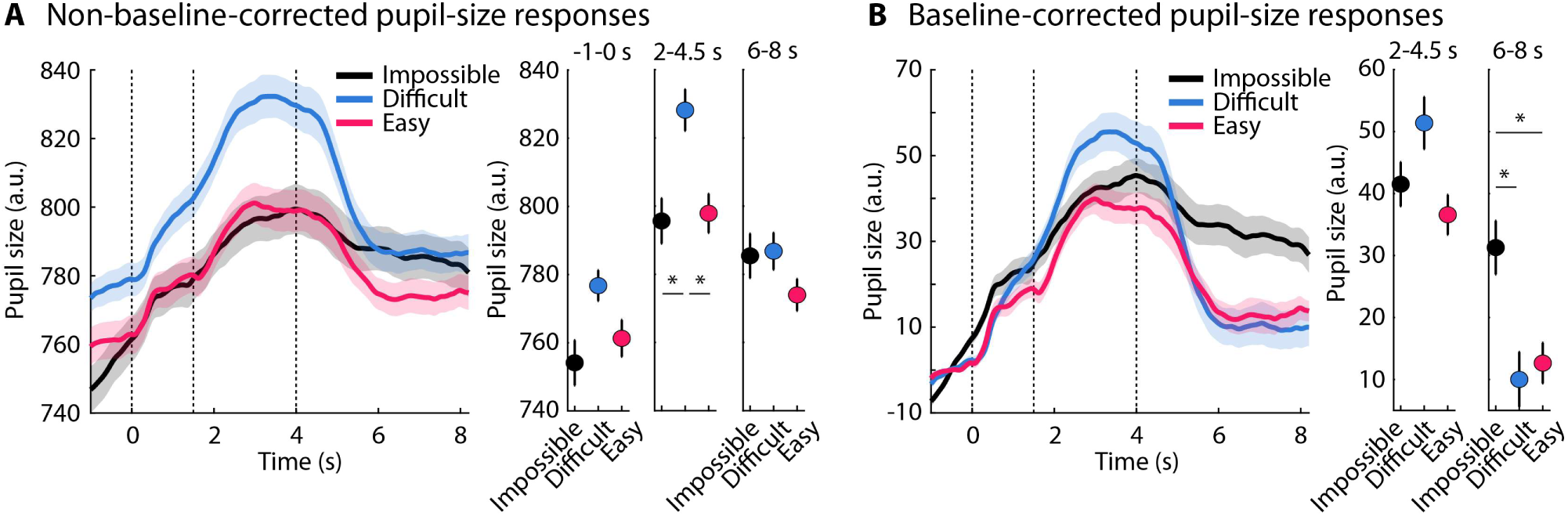
Pupil size results for Experiment 3. **A:** Non-baseline-corrected data, with time-courses shown on the left and mean pupil size for three time windows shown on the right. The dashed, vertical lines in the time-course plot mark the babble onset, sentence onset, and average sentence offset, from left to right respectively. Shaded areas around the time-courses and error bars reflect the standard error of the mean after removal of between-participant variance. *p_Holm_ ≤ 0.05; only significant effects are indicated. **B:** Same as for Panel A, for baseline-corrected pupil size.

Baseline-corrected pupil data also showed an SNR × time window interaction (F_2,8244_ = 14.992, p = 3.1 · 10^−7^, η_p_^2^ = 0.004). There were no condition differences for the sentence-response time window. For the post-sentence time window, similar to Experiment 2, pupil size was larger in the impossible relative to both the difficult condition (b = 21.97, 95% CI [7.05, 36.88], t_8244_ = 2.89, p_Holm_ = 0.023) and the easy condition (b = 19.00, 95% CI [5.51, 32.49], t_8244_ = 2.76, p_Holm_ = 0.029; Figure 5B).

#### Gaze dispersion

For non-baseline-corrected data, the SNR effects differed between time windows (interaction: F_4,12277_ = 50.728, p = 2 · 10^−49^, η_p_^2^ = 0.016). Gaze dispersion was lower throughout the trial – i.e., in all three time windows – for the impossible relative to the easy condition (for all, b ≥ 0.07, 95% CI [≥0.03, ≥0.11], t_12777_ ≥ 3.21, p_Holm_ ≤ 0.005). In the pre-noise and post-sentence time windows, gaze dispersion was also lower in the impossible compared to the difficult condition (for both, b ≥ 0.09, 95% CI [≥0.05, ≥0.14], t_12777_ ≥ 4.17, p_Holm_ < 0.001). In the sentence-response time window, gaze dispersion was lower for the difficult compared to both for the impossible condition (b = 0.06, 95% CI [0.01, 0.10,], t_12777_ = 2.57, p_Holm_ = 0.03) and the easy condition (b = 0.13, 95% CI [0.10, 0.15], t_12777_ = 8.81, p_Holm_ = 1.2 · 10^−17^; Figure 6A).

**Figure 6:**
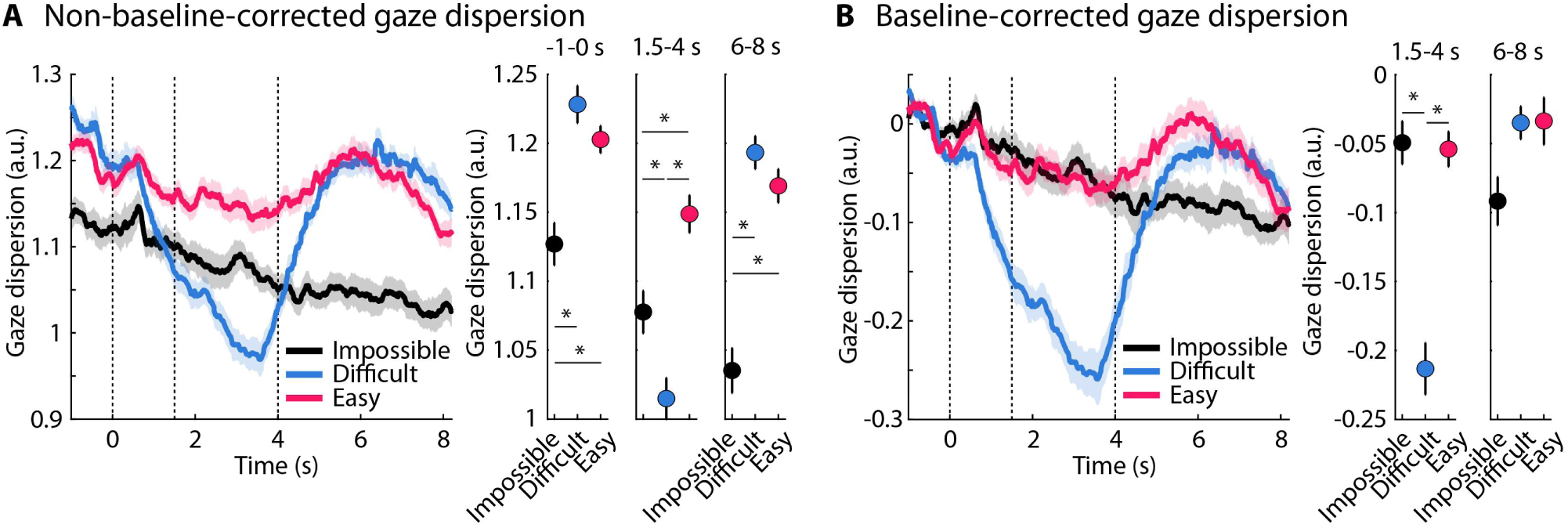
Gaze dispersion results for Experiment 3. **A:** Non-baseline-corrected data, with time-courses shown on the left and mean gaze dispersion for three time windows shown on the right. The dashed, vertical lines in the time-course plot mark the babble onset, sentence onset, and average sentence offset, from left to right respectively. Shaded areas around the time-courses and error bars reflect the standard error of the mean after removal of between-participant variance. *p_Holm_ ≤ 0.05; only significant effects are indicated. **B:** Same as for Panel A, for baseline-corrected gaze dispersion.

For baseline-corrected data (SNR × time window interaction: F_2,8070_ = 48.16, p = 1.6 · 10^−26^, η_p_^2^ = 0.012), gaze dispersion was lower for the difficult than the impossible (b = 0.15, 95% CI [0.10, 0.20], t_8070_ = 5.91, p_Holm_ = 1.8 · 10^−8^ ) and the easy condition (b = 0.14, 95% CI [0.10, 0.19], t_8070_ = 6.69, p_Holm_ = 1.5 · 10^−10^) in the sentence-response time window (1.5 to 4 s; Figure 6B). There were no other significant effects.

#### Summary

Results from Experiment 3 provide partial evidence for an inverted u-shaped effort profile. Pupil size was largest in the difficult condition, consistent with increased listening effort, but this pattern was only significant in the baseline-corrected data. This inverted u-shaped pattern may thus have involved both sustained arousal and sentence-related phasic effort. As in Experiment 2, baseline-corrected pupil size was larger in the impossible condition than in both the difficult and easy conditions after sentence offset.

The gaze-dispersion results showed a clear phasic reduction in eye movements during the processing of difficult-to-understand speech. Critically, the non-baseline-corrected gaze-dispersion data also showed a sustained reduction in eye movements throughout the trial in the impossible condition relative to the easy condition. This pattern indicates a persistent cognitive state, potentially reflecting a shift toward disengagement-related processes, such as sustained inward attention, when participants are not actively processing meaningful speech. We further explored this hypothesis in Experiment 4.

## Experiment 4

Gaze dispersion data from Experiment 3 suggest that participants may not have actively invested cognitive effort in listening. Instead, they may have disengaged and allowed their minds to wander. In the absence of engaging visual stimuli, mind wandering may have contributed to the reduced eye movements during impossible comprehension relative to attentive listening during easily comprehensible speech. This could explain why eye movements remained low throughout the trial, even before auditory stimulation. If this is the case, participants might move their eyes more when presented with something visually engaging, such as a short video clip. Reduced eye movements would then appear only during the presentation of a difficult-to-understand sentence. Experiment 4 was designed to test this hypothesis.

### Methods and materials

#### Participants

Twenty-six adults (median age: 24.5 years, age range: 19–33 years; 7 male, 18 female, 1 non-binary) participated in Experiment 4. Data from one additional participant were recorded but not analyzed because on more than 50% of trials, more than 50% of the physiological data were missing.

#### Stimulation and procedures

Experimental procedures were the same as in Experiment 3: no sentence was presented in the impossible condition, and participants were informed that the sentence would be absent. The only difference from Experiment 3 was that a short video clip was presented concurrently with the 8.5-s babble noise. To this end, a research assistant selected 180 neutral valence and arousal videos with a duration of at least 8.5 s from the Second Database of Emotional Videos from Ottawa (DEVO-2; Davidson et al., 2025). Videos longer than 8.5 s were clipped to match the duration of the babble noise. Video content comprised realistic everyday scenes and events, including natural landscapes, animals, people performing everyday actions, and object-centered scenes. Videos were randomly assigned to sentences, conditions, and participants.

### Results

#### Behavior

Performance in the semantic relatedness task was more accurate in the easy condition compared to both difficult and impossible conditions, and more accurate in the difficult than impossible condition (mean accuracy: 0.928, 0.719, and 0.515 for easy, difficult, and impossible, respectively; for all b ≥ 0.96, 95% CI [≥0.75, ≥1.16], OR ≥ 2.60, t_4677_ ≥ 9.28, p_Holm_ < 0.001; effect of SNR: F_2,4677_ = 138.96, p = 2.4 · 10^−59^). Performance in the impossible condition did not differ from chance level (b = 0.055, 95% CI [-0.048, 0.158], t_4677_ = 1.05, p = 0.295).

#### Pupil size

For non-baseline-corrected data, the SNR effect differed between time windows (SNR × time window interaction: F_4,12918_ = 4.736, p = 0.0008, η_p_^2^ = 0.001). For all time windows, the pupil size was greater in the difficult condition compared to both the impossible condition (for all, b ≥ 26.13, 95% CI [≥7.85, ≥44.41], t_12918_ ≥ 2.80, p_Holm_ ≤ 0.025). For the sentence-response and post-sentence time windows, the pupil size was also greater in the difficult condition than the easy condition (for both, b ≥ 30.14, 95% CI [≥12.82, ≥47.46], t_12918_ ≥ 3.41, p_Holm_ ≤ 0.004; Figure 7A).

**Figure 7:**
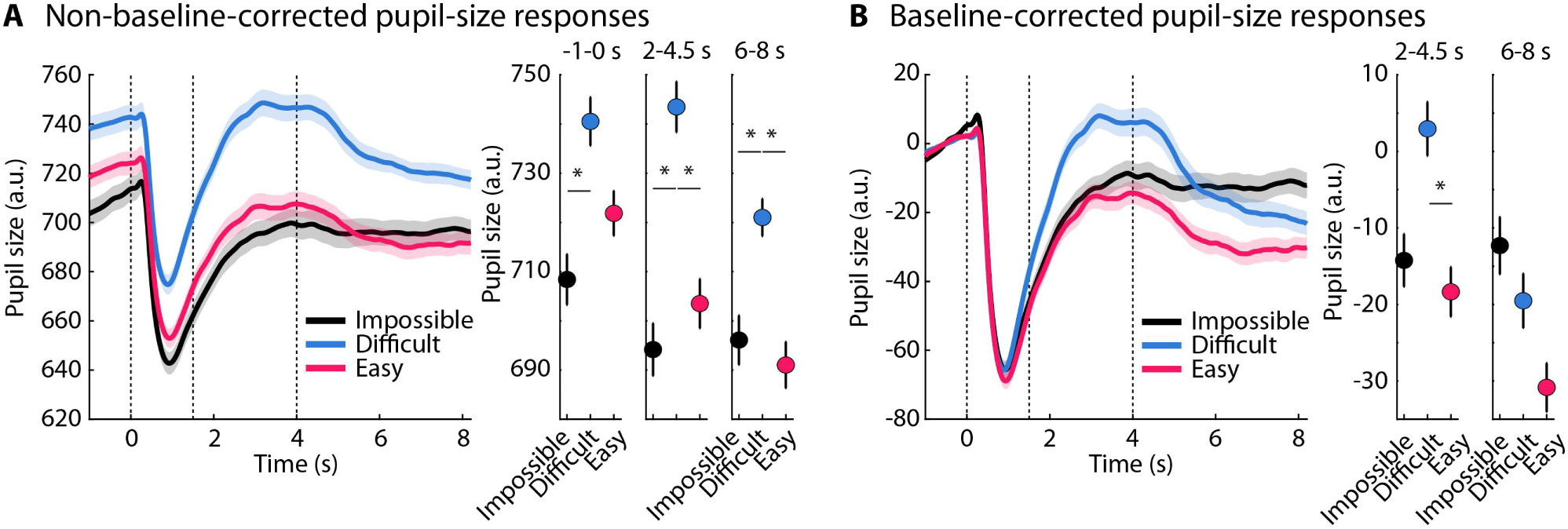
Pupil size results for Experiment 4. **A:** Non-baseline-corrected data, with time-courses shown on the left and mean pupil size for three time windows shown on the right. The video and babble noise started at time point zero and continued throughout the rest of a trial. The dashed, vertical lines in the time-course plot mark the babble onset, sentence onset, and average sentence offset, from left to right respectively. Shaded areas around the time-courses and error bars reflect the standard error of the mean after removal of between-participant variance. *p_Holm_ ≤ 0.05; only significant effects are indicated. **B:** Same as for Panel A, for baseline-corrected pupil size.

For baseline-corrected data (SNR × time window interaction: F_2,8612_ = 5.625, p = 0.0004, η_p_^2^ = 0.001), the pupil size was greater for the difficult condition compared to the easy condition sentence-response time window (b = 22.64, 95% CI [9.38, 35.90], t_8612_ = 3.35, p_Holm_ = 0.005; Figure 7B). Other effects were not significant after multiple-comparison correction (p_Holm_ > 0.05).

#### Gaze dispersion

Non-baseline-corrected data showed different SNR effects across time windows (interaction: F_4,12856_ = 18.501, p = 3.6 · 10^−15^, η_p_^2^ = 0.006). In the sentence-response time window (1.5 to 4 s), gaze dispersion was lower in the difficult condition than both the impossible condition (b = 0.07, 95% CI [0.04, 0.10], t_12856_ = 4.64, p_Holm_ = 2.8 · 10^−5^) and the easy condition (b = 0.12, 95% CI [0.09, 0.14], t_12856_ = 9.11, p_Holm_ = 8.8 · 10^−19^), and lower gaze dispersion in the impossible than in the easy condition (b = 0.05, 95% CI [0.01, 0.08], t_12856_ = 2.75, p_Holm_ = 0.030; Figure 8A). In the post-sentence time window (6 to 8 s), the gaze dispersion was lower for the impossible condition relative to both the difficult condition (b = 0.05, 95% CI [0.02, 0.08], t_12856_ = 3.03, p_Holm_ = 0.015) and the easy condition (b = 0.07, 95% CI [0.04, 0.10], t_12856_ = 4.28, p_Holm_ = 1.3 · 10^−4^). No other effects were significant.

**Figure 8:**
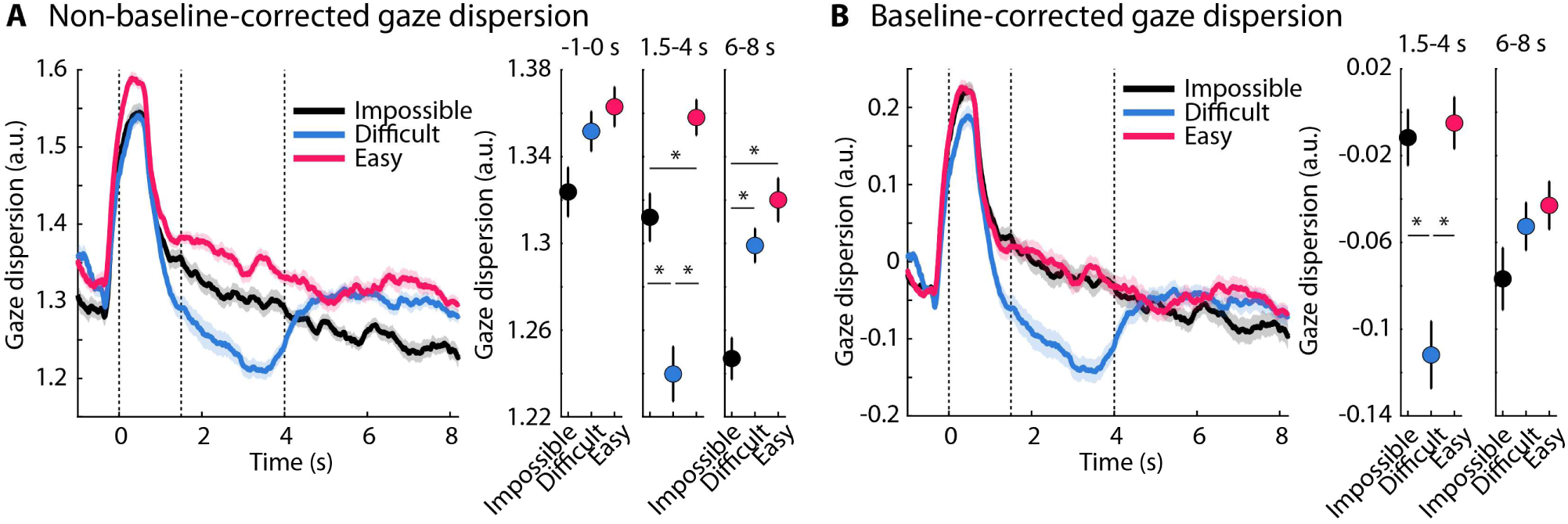
Gaze dispersion results for Experiment 4. **A:** Non-baseline-corrected data, with time-courses shown on the left and mean gaze dispersion for three time windows shown on the right. The video and babble noise started at time point zero and continued throughout the rest of a trial. The dashed, vertical lines in the time-course plot mark the babble onset, sentence onset, and average sentence offset, from left to right respectively. Shaded areas around the time-courses and error bars reflect the standard error of the mean after removal of between-participant variance. *p_Holm_ ≤ 0.05; only significant effects are indicated. **B:** Same as for Panel A, for baseline-corrected gaze dispersion.

For the baseline-corrected data (SNR × time window interaction: F_2,8496_ = 17.156, p = 3.7 · 10^−8^, η_p_^2^ = 0.001), gaze dispersion during the sentence-response time window was lower for the difficult condition compared to both the impossible condition (b = 0.09, 95% CI [0.04, 0.14], t_8496_ = 3.42, p_Holm_ = 0.003) and the easy condition (b = 0.10, 95% CI [0.06, 0.14], t_8496_ = 4.38, p_Holm_ = 7.2 · 10^−5^). No other effects were significant.

#### Summary

Experiment 4 tested whether engaging visual stimulation would reduce the sustained suppression of eye movements observed in the impossible condition. Pupil size again showed the expected effort-related pattern during sentence processing, with larger responses for difficult than for easy or impossible speech. Gaze dispersion was selectively reduced during difficult speech, consistent with reduced eye movements during effortful listening. Unlike Experiment 3, gaze dispersion did not differ significantly between conditions before auditory stimulation, suggesting that the videos encouraged visual exploration and reduced sustained pre-trial differences. However, non-baseline-corrected gaze dispersion remained lower for impossible than easy speech during and after sentence presentation. This pattern suggests that visual stimulation attenuated, but did not fully rescue, eye movements in the impossible condition, supporting the idea that gaze dispersion reflects both phasic listening demands and more sustained cognitive states. Baseline-corrected gaze dispersion showed the u-shaped function consistent with the expected effort profile (Yerkes and Dodson, 1908).

#### Comparison of difficulty effect between pupil size and gaze dispersion

In addition to examining whether pupil size and gaze dispersion each capture the full effort profile, we also asked whether the magnitude of the difficulty effect, defined as the difference between the easy and difficult conditions, differed between the two metrics. This comparison was motivated by two observations. First, the effect sizes in Experiments 2 and 3 appeared to be smaller for pupil size than for gaze dispersion. Second, previous work suggests that gaze dispersion may be able to capture differences between easy and difficult listening better than pupil size, at least in continuous listening paradigms (Herrmann et al., 2025). Comparing the magnitude of the difficulty effect across metrics may thus indicate under which experimental conditions each measure yields larger effects.

To compare the metrics, we analyzed the easy and difficult conditions separately for each experiment (sign-inverting gaze dispersion to mimic the direction of the pupil-size data). For each metric, we first calculated the mean and standard deviation across all participant values from the easy and difficult conditions. We then z-normalized each participant’s values by subtracting this mean and dividing by the standard deviation. This procedure preserved condition differences while allowing the difficulty effects to be compared across metrics and experiments. We then calculated the ‘difficulty effect’ as the difference between the difficult and easy conditions, separately for pupil size and gaze dispersion. For each experiment, a paired-samples t-test was used to compare the z-normalized difficulty effects between the two metrics.

The difficulty effect was larger for gaze dispersion than for pupil size in Experiments 2 and 3 (t_24_ = 2.409, p = 0.024, d = 0.482; t_25_ = 2.404, p = 0.024, d = 0.471, respectively), but not in Experiments 1 and 4 (t_23_ = 0.648, p = 0.523, d = 0.132; t_25_ = -0.317, p = 0.754, d = 0.062, respectively; Figure 9). This pattern suggests that gaze dispersion may capture differences between easy and difficult listening particularly well and better than pupil size during longer trials and in the absence of visual stimulation.

**Figure 9:**
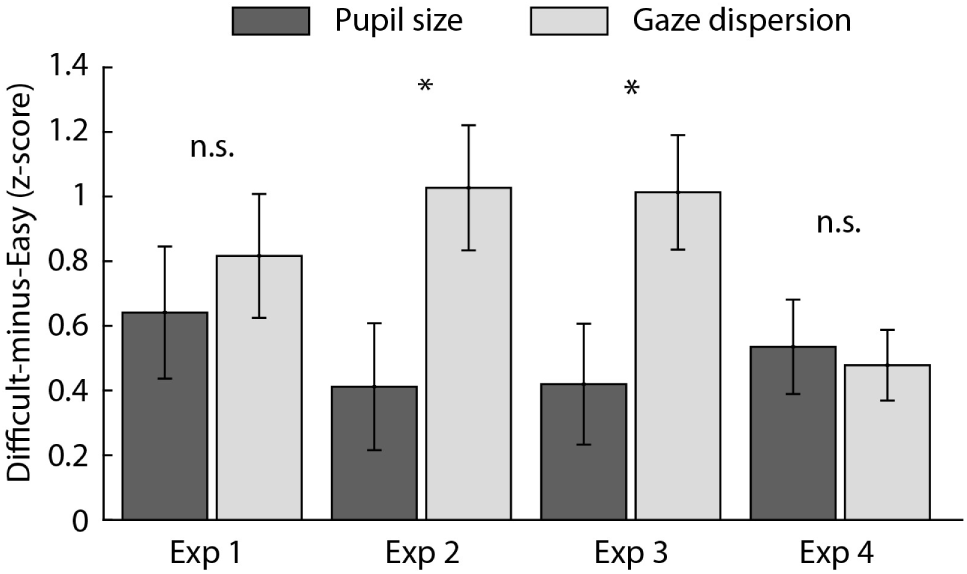
Comparison of the comprehension difficulty effect between pupil size and gaze dispersion. Bar graphs show the difference between the difficult and the easy condition, using z-normalized pupil size and gaze dispersion. Z-normalization enables comparing the difficulty effect across metrics (pupil size, gaze dispersion) and experiments. Error bars reflect the standard error of the mean. *p ≤ 0.05, n.s. – not significant.

## Discussion

The current study examined how pupil size and eye movements track understanding speech that is easy, difficult, and impossible to follow. Across four experiments, pupil size and gaze dispersion showed related but dissociable profiles. Pupil size generally followed the expected effort pattern, with larger responses during difficult than easy or impossible listening. Gaze dispersion also decreased during difficult speech processing, consistent with effortful listening, but showed sustained reductions when comprehension was impossible. Together, the findings fit with an arousal-related interpretation of pupil size, while suggesting that gaze dispersion reflects both attentional engagement during active listening and disengagement-related states when meaningful listening is no longer possible.

### Pupil size reflects listening effort

Across experiments, pupil size was generally larger for difficult but still intelligible speech than for easy or impossible speech, showing the expected inverted u-shape (Yerkes and Dodson, 1908; Zekveld and Kramer, 2014; Ohlenforst et al., 2017b; Ohlenforst et al., 2018; Wendt et al., 2018; Herrmann and Ryan, 2024) and reinforcing the utility of pupillometry as an index of listening effort. The effort profile was, however, not equally strong across analyses, and the difficult-versus-easy and difficult-versus-impossible differences were weaker or did not survive correction for multiple comparisons in some cases, especially for baseline-corrected data (Figures 3 and 5). Pupil responses in the present paradigm thus appear to reflect a combination of sustained processes and phasic, sentence-evoked effort. Because the signal-to-noise ratio conditions were cued and blocked, participants could anticipate whether a trial would be easy, difficult, or impossible, and may have prepared for difficult listening by increasing arousal, and in turn pupil size (Mathôt, 2018; Joshi and Gold, 2020; Strauch et al., 2022), before and throughout a trial, and this appeared to contribute to the inverted u-shaped pattern in the current study.

A further notable finding was the larger pupil response in the impossible condition after sentence offset, particularly in baseline-corrected analyses in Experiments 2 and 3. Its late timing, when no speech could occur, makes listening effort an unlikely explanation. One possibility is arousal related to the upcoming response, because participants had to guess whether the probe word was related to a sentence they could not understand or that was absent, and this uncertainty could have increased arousal before the probe. Alternatively, the effect may reflect a relative decrease in the easy and difficult conditions, because listeners who understood the sentence could already resolve the judgment while the sentence unfolded. Both accounts locate the late response in processes surrounding the behavioral response rather than in listening. Previous studies using signal-to-noise ratios at which participants appeared disengaged have not reported pupil time courses (Zekveld and Kramer, 2014; Ohlenforst et al., 2017b; Ohlenforst et al., 2018), used trials too short to reveal such late responses (Herrmann and Ryan, 2024), or showed a similar increase for impossible conditions without analyzing it statistically (Wendt et al., 2018). This pattern may thus need be considered when interpreting pupil responses in paradigms that include disengagement-inducing conditions.

### Eye movements reflect attentional engagement during active listening

In all four experiments, we observed a phasic reduction in gaze dispersion for the difficult compared to the easy condition during the sentence-listening period of a trial. This is consistent with recent works showing reduced eye movements when speech is masked by background noise (Contadini-Wright et al., 2023; Cui and Herrmann, 2023; Herrmann and Ryan, 2024; Mariotti et al., 2026), and during high compared to low memory load (Lipton et al., 1980; Hutton and Tegally, 2005; Kosch et al., 2018; Walter and Bex, 2021). The current work shows that this reduction in eye movements occurs even while a video is presented (Experiment 4).

Critically, gaze dispersion did not simply mirror pupil size. During sentence listening the two measures agreed, with larger pupil size and lower gaze dispersion for difficult than easy speech, in both non-baseline-corrected and baseline-corrected data. They differed in how much of this effect was carried by sustained differences preceding the sentence. For pupil size, the difficult-versus-easy effect was weaker or absent after baseline correction (Experiments 2 and 3), and in Experiment 4 pupil size was already larger in the difficult than the impossible condition before noise onset, indicating an anticipatory contribution. For gaze dispersion, the reduction during difficult listening was similar with and without baseline correction, and no experiment showed easy-versus-difficult differences in the pre-noise or post-sentence windows. Reduced eye movements during difficult listening thus appear tied to the period during which a sentence was present, whereas the pupil-size effect also reflected a sustained preceding state. This is in line with the proposal that pre-stimulus baseline pupil size reflects anticipatory arousal and preparatory processing related to task difficulty, whereas the evoked response reflects effort during listening (Ayasse and Wingfield, 2020; Relaño-Iborra et al., 2022; Seropian et al., 2022), and with suggestions that pupil size and eye movements index partly overlapping but not identical mechanisms during listening (Herrmann and Ryan, 2024; Huviyetli and Chait, 2026; Liu and Chait, 2026).

Pupil size is closely linked to arousal systems, particularly the locus coeruleus– norepinephrine system (Bradley et al., 2008; Mathôt, 2018; Wang et al., 2018; Joshi and Gold, 2020; Ross and Van Bockstaele, 2021; Burlingham et al., 2022; Strauch et al., 2022; Torres et al., 2025). Eye movements, including small fixational movements and broader gaze exploration, are governed by partly overlapping but distinct oculomotor and attentional networks, including cortical and subcortical structures involved in orienting, visual exploration, and cognitive control (Sparks, 2002; Pierrot-Deseilligny et al., 2004; Lencer and Trillenberg, 2008; Pierce et al., 2019). Reduced gaze dispersion during difficult listening may reflect a shift in attentional engagement, such that listeners may reduce visual exploration to prioritize auditory processing or to minimize irrelevant visual input while trying to understand masked speech.

### Sustained eye-movement reductions may reflect disengagement or inward attention

The most important departure from a simple effort account was the sustained reduction in gaze dispersion during impossible listening. We had expected a u-shaped profile (Yerkes and Dodson, 1908), with similarly high gaze dispersion (indexing low effect) for easy and impossible speech, assuming that listeners have little motivation to attempt comprehension when additional effort cannot improve listening success (Brehm and Self, 1989; Botvinick and Braver, 2015; Herrmann and Johnsrude, 2020). This pattern emerged in baseline-corrected analyses, particularly in Experiments 3 and 4 (Figures 6B and 8B), but non-baseline-corrected data in Experiments 2 and 3 showed gaze dispersion to remain lower throughout the trial, including the pre-noise period, in the impossible condition (Figures 4A and 6A). Because the effect occurred without visual stimulation, it cannot be attributed to suppressed exploration of the moving-dot display used previously (Herrmann and Ryan, 2024), and because neither extending the post-sentence interval nor removing the sentence restored eye movements, it cannot reflect a continued search for an inaudible sentence that would end once listeners inferred the sentence was over. Reduced eye movements thus do not always index attentional engagement or listening effort, and the effect does not require attempts to extract speech from noise.

We suggest that sustained eye-movement reductions during impossible listening reflect a broader cognitive state, such as disengagement, inward attention, or mind wandering (Smallwood and Schooler, 2006; Smallwood and Andrews-Hanna, 2013; Westgate and Wilson, 2018). When no useful speech information can be extracted and little of interest is on the screen, participants may withdraw from both auditory and visual engagement, reducing visual exploration. This is consistent with reports of reduced eye movements during mind wandering in reading (Reichle et al., 2010; Uzzaman and Joordens, 2011; Mézière et al., 2026; but see Faber et al., 2020) and during internally compared to externally focused attention (Benedek et al., 2017; Walcher et al., 2017). Reduced eye movements may thus arise from two states, focused engagement during difficult listening and disengaged inward focus during impossible listening, whose distinction may depend on timing, baseline correction, and task context. Notably, gaze dispersion was lower and more closely confined to the sentence-listening period in the difficult condition than in the impossible condition (Figures 2A, 6A, 8A). The fixed trial timing further means that comprehension in the easy condition was typically complete well before a trial ended, such that participants could also have disengaged during the post-sentence period. Gaze dispersion in the easy condition did not, however, decrease after sentence offset and remained higher than in the impossible condition (Figures 4A, 6A), suggesting that completed comprehension is not sufficient to induce a sustained reduction in eye movements, and that advance knowledge that nothing meaningful could be understood matters more.

Experiment 4 tested the disengagement interpretation by presenting videos during the trial to encourage visual engagement. The manipulation reduced some of the sustained pre-trial differences in gaze dispersion, suggesting that visual engagement can partly counteract disengagement-related reductions in eye movements, but reduced gaze dispersion in the impossible condition was not fully eliminated (Figure 8A). One possibility is that the videos were not sufficiently engaging to maintain external attention throughout blocked impossible trials, as the clips were neutral in valence and arousal and many contained relatively localized visual action (Davidson et al., 2025). Another is that knowing no sentence was present promoted disengagement from the trial as a whole, even with visual input. Future work could test this using more engaging visual scenes, trial-by-trial measures of mind wandering, or conditions in which participants have an explicit visual task during impossible listening.

A limitation of the present work is that our design does not capture the dynamics of disengagement. Blocking and cueing the signal-to-noise ratio conditions was intentional to facilitate disengagement in the impossible condition (Herrmann and Ryan, 2024), but it did not create the everyday situation in which a listener withdraws effort only after gradually realizing that comprehension is impossible. We further obtained no subjective report of invested effort or of sentence detection, because probing sentence presence would have rendered the acoustic signal task-relevant in the very condition in which participants were expected to abandon listening. The similar results of Experiments 2 and 3 suggest that sentence detection was not critical, although future work should complement physiological measures with direct reports, ideally when listening difficulty is not known in advance. The sustained reduction in gaze dispersion could further reflect preparation for a decision under complete uncertainty rather than disengagement, although gaze dispersion was already reduced during the pre-noise window and participants in Experiments 3 and 4 knew that no sentence would be presented. Such a contribution cannot be fully excluded, and future work could omit the semantic-relatedness judgment in the impossible condition to test whether both measures return to baseline.

### Implications for using eye movements as a measure of listening effort

The present findings have two implications for using eye movements as an index of listening effort. First, gaze dispersion is a reliable marker of active listening engagement (Contadini-Wright et al., 2023; Cui et al., 2025). Across experiments, gaze dispersion decreased during difficult speech processing, specifically while a sentence was present. The size of this effect relative to pupil size varied across experiments. For longer trials and in the absence of visual stimulation, the difficult-to-easy effect was larger for gaze dispersion than for pupil size (Experiments 2 and 3; Figure 9), consistent with reports that pupil-size effects can be reduced while gaze-dispersion effects remain robust during continued story listening (Cui and Herrmann, 2023; Herrmann et al., 2025; Widmann et al., 2025). Assessing gaze dispersion may thus be useful in contexts in which pupillometry is difficult to interpret, such as paradigms with less controlled luminance, more naturalistic viewing, or long continuous listening periods. Which measure is more useful in a given study likely also depends on practical constraints such as the number of trials, the population being tested, and other design choices.

Second, reduced eye movements (lower gaze dispersion) should not automatically be interpreted as increased listening effort, a caveat which appears to apply to increased pupil size as well (Figures 3B, 5B, 7B). The sustained reduction in gaze dispersion during impossible listening shows that similar eye-movement patterns can arise from different cognitive states. During difficult listening, reduced gaze dispersion may reflect attentional engagement with speech. During impossible listening, a sustained reduction in gaze dispersion may instead reflect disengagement, inward attention, or reduced exploration of the external environment. This means that gaze dispersion is most informative when interpreted together with task structure, behavioral performance, the time course of the effects, and, where possible, other physiological measures such as pupil size. The partial dissociation between pupil size and gaze dispersion suggests the two measures could provide complementary information (cf. Contadini-Wright et al., 2023; Herrmann and Ryan, 2024; Huviyetli and Chait, 2026; Liu and Chait, 2026): pupil size may be more closely aligned with attention allocation that is accompanied by arousal and effort investment, whereas gaze dispersion may also capture shifts of attention between external sensory processing, visual exploration, and internally oriented states that do not require effort investment. Pupil size is, however, also sensitive to attentional allocation and to internally directed states (Koelewijn et al., 2015; Unsworth and Robison, 2016; Zekveld et al., 2018), and the present data do not isolate the mechanisms underlying either measure.

## Conclusion

Across four experiments, pupil size and gaze dispersion provided complementary indices of listening-related cognition. Pupil size generally followed the expected inverted u-shape effort profile, increasing most when speech was difficult but still understandable. Eye movements – as assessed using gaze dispersion – showed a robust phasic reduction during difficult listening, consistent with increased attentional engagement during active speech processing. However, gaze dispersion also remained low during impossible listening in non-baseline-corrected data, suggesting that reduced eye movements can reflect disengagement or inward attention rather than effort alone. These findings suggest that eye movements are a sensitive marker of attentional engagement during listening, but their interpretation depends on context. Combining pupil size and eye movements may therefore provide a richer account of how listeners allocate effort, attention, and arousal across changing listening demands.

## Data availability

Data will be available upon reasonable request to the corresponding author.

## Author Contributions

**BH:** Conceptualization, methodology, validation, formal analysis, investigation, data curation, writing - original draft, writing - review & editing, visualization, supervision, project management, funding acquisition. **LKF:** Conceptualization, methodology, investigation, writing - review & editing. **PRP:** Investigation, data curation, writing - review & editing. **ISJ:** Conceptualization, methodology, investigation, writing - review & editing. **JDR:** Conceptualization, methodology, investigation, writing - review & editing.

## Acknowledgements

The research was supported by the Canada Research Chair program (CRC-2023-00383), the Natural Sciences and Engineering Research Council of Canada (NSERC Discovery Grant: RGPIN-2021-02602), and the Canadian Institutes of Health Research (CIHR: 186236). We thank Sarah Bobbitt, Lisa D’Souza, and Rojin Firooz for their help during data collection.

## Statements and Declarations

The authors have no conflicts or competing interests.

